# Conformational dynamics of Cas9 governing DNA cleavage revealed by single molecule FRET

**DOI:** 10.1101/167627

**Authors:** Mengyi Yang, Sijia Peng, Ruirui Sun, Jingdi Lin, Nan Wang, Chunlai Chen

**Affiliations:** School of Life Sciences; Tsinghua-Peking Joint Center for Life Sciences; Beijing Advanced Innovation Center for Structural Biology, Tsinghua University, Beijing, China.

## Abstract

Off-target binding and cleavage by Cas9 pose as major challenges in its applications. How conformational dynamics of Cas9 governs its nuclease activity under on- and off-target conditions remains largely unknown. Here, using intra-molecular single molecule fluorescence resonance energy transfer measurements, we revealed that Cas9 in apo, sgRNA-bound, and dsDNA/sgRNA-bound forms all spontaneously transits between three major conformational states, mainly reflecting significant conformational mobility of the catalytic HNH domain. We furthermore uncovered a surprising long-range allosteric communication between the HNH domain and RNA/DNA heteroduplex at the PAM-distal end to ensure correct positioning of the catalytic site, which demonstrated a unique proofreading mechanism served as the last checkpoint before DNA cleavage. Several Cas9 residues were likely to mediate the allosteric communication and proofreading step. Modulating interactions between Cas9 and heteroduplex at the distal end by introducing mutations on these sites provides an alternative route to improve and optimize the CRISPR/Cas9 toolbox.

## Introduction

CRISPR (Clustered Regularly Interspaced Short Palindromic Repeats)-associated protein Cas9 is a multi-domain DNA endonuclease that functions as a part of bacterial adaptive immune system to protect the host against the invasion of virus or foreign plasmids (Barrangou et al., 2007; Wiedenheft et al., 2012; Wright et al., 2016). Guided by a dual-tracrRNA:crRNA or a chimeric single-guide RNA (sgRNA), Cas9 targets 20 base pair (bp) complementary sequences for site-specific double-strand DNA breaks (Jinek et al., 2012; Jinek et al., 2014). Various structural, biochemical, and single molecule studies have revealed that Cas9 firstly samples and recognizes protospacer adjacent motif (PAM) on the DNA sequence. Complementarity between RNA guide sequence and target DNA strand, especially for “seed” sequence, leads to partial RNA-DNA heteroduplex formation in the PAM-proximal region and extended to the distal end to form a “R-loop”, which positions the catalytic active site of Cas9 adjacent to 3 bp upstream of the PAM to cleave target DNA (Jiang et al., 2016; Jinek et al., 2012; O’Connell et al., 2014; Singh et al., 2016; Sternberg et al., 2015; Sternberg et al., 2014; Szczelkun et al., 2014; Wright et al., 2015). The simplicity and specificity of programmable DNA recognition and cleavage by CRISPR/Cas9 enable applications of precision genome editing and manipulation across many areas of biology (Barrangou and Doudna, 2016; Cong et al., 2013; Hsu et al., 2014).

Cas9 has been shown to recognize and cleave DNA target bearing imperfect complementarity towards RNA guide sequence, particularly when mismatches are present at the PAM-distal end (Fu et al., 2013; Hsu et al., 2013; Lim et al., 2016; Sternberg et al., 2015). Such off-target effect poses as one of the major challenges in its applications. Several methods have been reported to improve the specificity of Cas9, including applying with truncated sgRNA and engineered Cas9 variants with attenuated interactions with DNA strands (Fu et al., 2014; Kleinstiver et al., 2016; Slaymaker et al., 2016). Using ensemble FRET measurements, Sternberg and coworkers have shown that the catalytic HNH domain of Cas9 is highly dynamic and its conformational state correlates with DNA cleavage activity (Sternberg et al., 2015). However, the conformational dynamics and molecular mechanisms of Cas9 governing its nuclease activity under on- and off-target conditions remain largely unresolved.

Here, we applied single molecule fluorescence resonance energy transfer (smFRET) to directly characterize conformational dynamics of individual *Streptococcus pyogenes* Cas9. We showed that Cas9 spontaneously fluctuates among three conformational states accompanied by remarkable movements of the catalytic HNH domain. A compact closed conformational state, which should closely resemble the cleavage-competent state of Cas9, is least populated under all experimental conditions. In addition, we examined how the dynamics of Cas9 was affected by introducing mismatches between sgRNA and target DNA strand, mismatches between target and non-target DNA strands, and mutations on Cas9 residues. Our results suggested that Cas9 utilizes a proofreading mechanism through a long-range allosteric communication between the PAM-distal end and its catalytic HNH domain. Interactions between several Cas9 residues and RNA/DNA heteroduplex at the distal end trigger Cas9 to transiently sample its “cleavage-competent” closed state to mediate DNA cleavage, whereas 4 or more mismatches between sgRNA and DNA cause incorrect positioning of the HNH domain and a transiently-formed “cleavage-impaired” closed state. Our findings provide fundamental molecular insights to understand how conformational dynamics of Cas9 and interactions between Cas9 and RNA/DNA heteroduplex govern its nuclease activity and target specificity. Finally, we propose an alternative route to improve Cas9 specificity by mutating its residues involved in the proofreading step.

## Results

### Design for smFRET

Crystal structures have shown that Cas9 adopts quite different conformations in its Apo, sgRNA-bound, incomplete DNA/sgRNA-bound, and double-stranded DNA(dsDNA)/sgRNA-bound forms (Fig. 1a). The conformation of catalytic HNH domain is relative conserved and Cas9 exhibits an “open” conformation in apo and sgRNA-bound forms (Jiang et al., 2015; Jinek et al., 2014). Binding with an incomplete DNA displays an “intermediate” Cas9 conformation with a remarkable movement of HNH domain towards the RNA/DNA heteroduplex (Anders et al., 2014; Nishimasu et al., 2014). In the dsDNA/sgRNA-bound form, Cas9 adopts a more compact (“closed”) conformation accompanied with a profound ~180°rotation and a further ~20 Å movement of the HNH nuclease domain towards the DNA cleavage sites (Jiang et al., 2016). We chose two intra-molecular FRET pairs, S355-S867 and S867-N1054 (Sternberg et al., 2015), which displayed distinguishable changes of relative distances among open, intermediate, and closed conformational states of Cas9 (Fig. 1a). By introducing cysteine at these sites on a cysteine free variants, we generated Cas9_HNH1_ (C80S/S355C/C574S/S867C) and Cas9_HNH2_ (C80S/C574S/S867C/N1054C) for Cy3 and Cy5 labeling. Labeled Cas9_HNH1_ and Cas9_HNH2_ had similar nuclease activities as wild type Cas9 (Fig. S1).

**Fig. 1.**
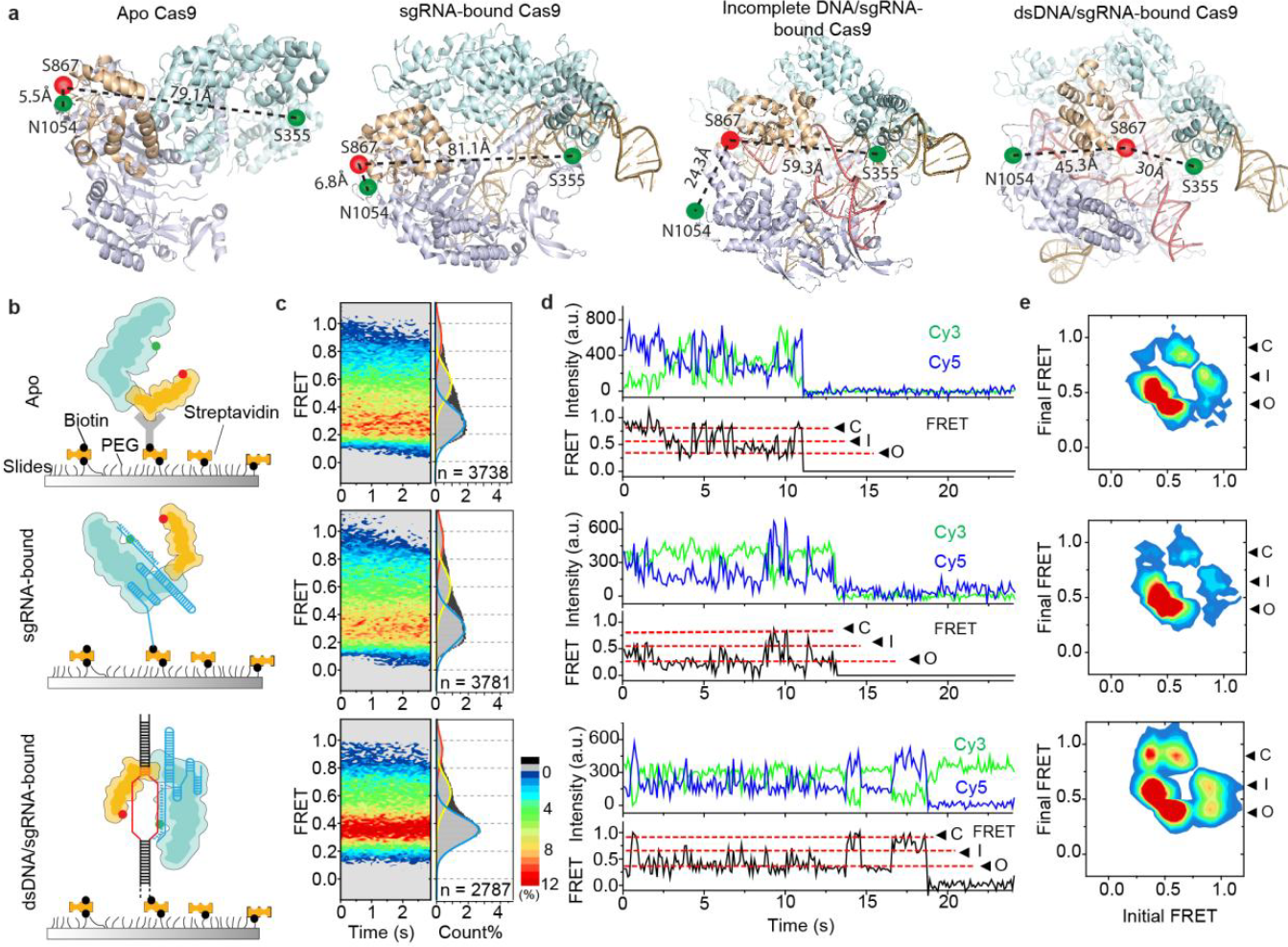
Conformational flexibility of Cas9. **a**, Structures of Cas9 displaying fluorophore labeling sites and distances between FRET pairs of Cas9_HNH1_(S355/S867) and Cas9_HNH2_(S867/N1054) in apo (pdb: 4CMP), sgRNA-bound (pdb: 4ZTO), incomplete DNA/sgRNA-bound (pdb: 4UN3), and dsDNA/sgRNA-bound forms (pdb: 5F9R). HNH domain and α-helical lobe are shown in wheat and palecyan, respectively. The rest domains are all shown in bluewhite. **b**, Cartoon demonstrating how different forms of Cas9 were attached on microscope slides for smFRET imaging. HNH domain and α-helical lobe of Cas9 are shown in wheat and cyan, respectively. Target DNA sequences are shown in red with the PAM in yellow, and sgRNA is colored in blue. **c**, FRET efficiency distributions for the apo (upper), sgRNA-bound (middle) and dsDNA/sgRNA-bound (lower) Cas9_HNH1_. Population contour plots (left) are normalized with counts number and are color-coded from navy (lowest) to red (highest population) with the color scale shown beside the graphs. The cumulative population histogram (right) displays three distinct FRET states (red, yellow and blue lines were Gaussian fitting of each FRET population, representing closed (C), intermediate (I), and open (O) states, respectively). **d**, Corresponding representative single molecule FRET traces for the apo, sgRNA-bound and dsDNA/sgRNA-bound Cas9_HNH1_ displaying spontaneous transitions between three distinct FRET states (C, I, and O). Sudden decreases of fluorescence signals to around 0 in single molecule traces were caused by photobleaching of Cy3 or Cy5 fluorophores. **e**, Corresponding transition density plots showing the frequency of transitions between states, in which initial and final FRET values for each transition are accumulated into two-dimensional histograms. Color scale is from blue (lowest frequency) to red (highest frequency).

### Conformational dynamics of Cas9 in steady state

Apo, sgRNA-bound, and dsDNA/sgRNA-bound Cy3/Cy5 labeled Cas9_HNH1_ and Cas9_HNH2_ were immobilized on PEG passivated microscope glass slide via biotinylated his-tag antibody, biotinylated sgRNA, and biotinylated dsDNA, respectively (Fig. 1b and Table S1). Objective-based total internal reflection fluorescence (TIRF) microscope was used to capture smFRET trajectories of individual Cas9 molecules under equilibrium conditions (steady state), which enabled us to track conformational dynamics of Cas9 catalytic HNH domain by monitoring the relative distance changes between Cy3 and Cy5 labeling sites. Strikingly, we discovered that both Cas9_HNH1_ and Cas9HNH2 exhibited and spontaneously transited among three distinct FRET states in all different forms (Fig. 1c-e, and S2). FRET values of Cas9_HNH1_ and Cas9_HNH2_ were in line with the relative distances between labeling sites (Tables S2 and S3), allowing us to assign the low FRET Cas9_HNH1_ and high FRET Cas9_HNH2_ to apo or sgRNA-bound like state (termed as “open state”), the medium FRET Cas9_HNH1_ and Cas9_HNH2_ to the incomplete DNA/sgRNA-bound like state (termed as “intermediate state”), and the high FRET Cas9_HNH1_ and low FRET Cas9_HNH2_ as the dsDNA/sgRNA-bound like state (termed as “closed state”). The open state was dominant in most cases, even when Cas9 bound on fully matched DNA. On the other hand, the closed state, which should closely resemble the cleavage-competent state of Cas9, was the least populated state in all cases. In all, smFRET measurements unambiguously captured the intrinsic structural flexibility of Cas9 conformation in real-time, which has been suggested by structural studies and molecular dynamic simulations (Anders et al., 2014; Jiang et al., 2016; Jiang et al., 2015; Jinek et al., 2014; Nishimasu et al., 2014; Palermo et al., 2016).

### Conformational dynamics of Cas9 with PAM-distal end mismatches between sgRNA and DNA

Next, we introduced 1-4 bp mismatches at the PAM-distal end to examine how off-target modulates conformational dynamics of Cas9 (Fig 2a). Similar to targeting perfectly matched DNA, both Cas9_HNH1_ and Cas9_HNH2_ exhibited three fluctuating states on all DNAs containing mismatches (Fig. 2b, S3, and S4). However, proportion of the open state increased ~15% when there were mismatches between target DNA and sgRNA (Fig 2c, Tables S2 and S3). To quantify thermodynamic and dynamic properties of three Cas9 conformational states, individual single molecule FRET trajectories were analyzed by a Hidden Markov Model based software [HaMMy (McKinney et al., 2006)] to assign each time point to one of three major FRET states (Fig. S3 and S4). More than 80% of individual molecules showed spontaneous transitions between two or more FRET states, from which dwell times of each state and transition rates among three Cas9 states were extracted (details in SI, Table S2 and S3). As the major population, the open state was defined as the ground state, whose free energy was set as 0. Free energies (Δ*G*) of the intermediate and closed states (Fig. 2d and S5a) were calculated using equation Δ*G* = –*k*_B_*T*ln*K*, in which *k*_B_ is the Boltzmann constant, *T* is the temperature, and *K* is the equilibrium constant calculated as the ratio of forward and backward transition rates between two states. Clearly, mismatches between sgRNA and target DNA destabilized both intermediate and closed states by 0.5-1.5 *k*_B_*T*. In addition, these mismatches also caused extra energy barriers on transition pathways which led to 3-6 folds slower effective transition rates from open and intermediate states to the closed state (Fig. 2e and S5b). Therefore, both thermodynamic and dynamic parameters supported the fact that mismatches destabilized the intermediate and closed states and increased energy barriers so that the closed Cas9 state, a potential cleavage-competent state, is harder to be accessed.

**Fig. 2.**
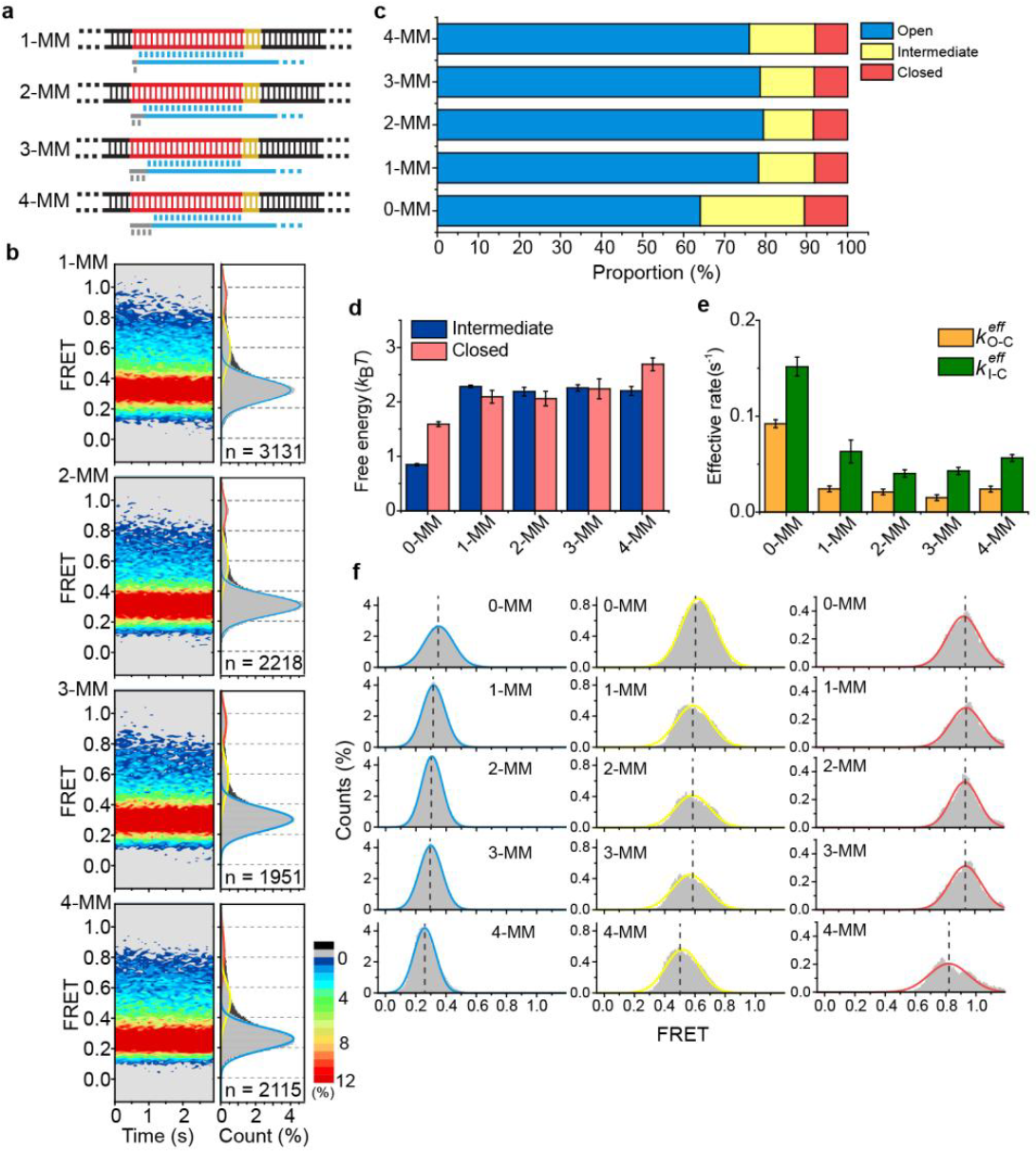
Conformational dynamics of Cas9_HNH1_ in the presence of PAM-distal end mismatches in steady state. **a**, Mismatches (MM) were introduced at the PAM-distal end between target DNA and sgRNA. Target sequences are shown in red with the PAM in yellow. Matched and mismatched bases of sgRNA towards target DNA are colored in blue and grey, respectively. **b**, FRET contour plots (left) and cumulative histograms (right) of sgRNA/Cas9_HNH1_ bound on DNA substrates containing 1-4 MM captured in steady state. **c**, Population percentage of three Cas9 conformational states. Proportion of the open state increased ~15% when there were mismatches between target DNA and sgRNA. **d**, Free energies of the intermediate and closed states of dsDNA/sgRNA-bound Cas9_HNH1_ in steady state. The open state is defined as the ground state whose free energy is 0. **e**, Effective transition rates of dsDNA/sgRNA-bound Cas9_HNH1_ from open and intermediate states to the closed state, 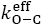 and 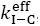, respectively. 0-MM stands for fully complementarity between DNA and sgRNA. **f**, FRET distributions of the closed (red), intermediate (yellow), and open (blue) Cas9_HNH1_ states in the presence of 0~4-MM DNA. FRET values of each time point were assigned by HaMMy (McKinney et al., 2006) to be classified into three different FRET states to generate these distributions. Colored solid curves were Gaussian fitting. Center of FRET peaks (marked by dash line) in the presence of 4-MM were significantly lower than other conditions.

As our biochemical assay (Fig. S3a) and previous studies have assessed (Fu et al., 2014; Sternberg et al., 2015), 4 or more mismatches at the PAM-distal end severely hindered or even abolished cleavage activity of Cas9. However, our steady-state smFRET measurements still captured the closed state in the presence of 4 mismatches. Thermodynamic stability and access rates of the closed state in the presence of 4 mismatches were almost the same as the ones with 1-3 mismatches (Fig. 2d-e and S5). Based on the assignment from HaMMy(McKinney et al., 2006), FRET values of each time point were classified into three groups to generate FRET distributions of the closed, intermediate, and open Cas9_HNH1_ states (Fig. 2f). Although full widths at half maximum (FWHM) of the peak were in the range of 0.17-0.30 for these FRET distributions, their peak positions were determined with high precision through Gaussian fitting (Fig. 2f, Table S2 and S3), which is the same approach used by several super-resolution imaging methods to accurately locate fluorophore position through two-dimensional Gaussian fitting (Forkey et al., 2003; Rust et al., 2006; Shroff et al., 2007). We noticed that the FRET values of the closed Cas9_HNH1_ state with 1-3 mismatches were identical to that without mismatches (FRET efficiency *E* = ~0.92 ± 0.01, Fig. 2f, Table S2), whereas 4 mismatches caused a significant decrease of the high FRET value to *E* = 0.81 ± 0.01. Similar results were found with 8 mismatches at the distal end (Table S2). Such significant FRET decrease corresponded to ~ 5 Å increase of the relative distance between labeling sites S355C and S867C, suggesting that the HNH nuclease domain might slightly move away from its catalytic site, which subsequently hindered or abolished its nuclease activity. Our results demonstrated a remarkable local conformational plasticity of the HNH domain even when Cas9 adopts a closed state, which has also been proposed by molecular dynamic simulations (Palermo et al., 2016). Such conformational plasticity should be highly localized, which was not sensed by the FRET pairs of Cas9_HNH2_ (Table S3). In summary, in addition to their ability to destabilize the closed state, 4 or more mismatches allosterically drived the local conformation of HNH domain away from its active form, implying that the closed state captured by our smFRET did not always represent the cleavage-competent state. Base pairing between DNA and sgRNA at the distal end allosterically triggers a local reposition of HNH toward its catalytic site, switching the closed state from a cleavage-impaired state (Cas9_HNH1_, *E* = 0.81 ±0.01) to a cleavage-competent state (Cas9_HNH1_, *E* = ~0.92 ±0.01).

### Conformational dynamics of Cas9 in pre-steady state

To fully understand how cleavage is governed by the conformational dynamics of Cas9 upon DNA binding, we captured the smFRET of Cas9 in pre-steady state by imaging 10 s prior to delivery of labeled Cas9_HNH1_ to surface immobilized DNA strands and examined the evolution of smFRET of Cas9_HNH1_ from the moment its bound to DNA (Fig. 3 and S6a). Cas9_HNH1_ mainly presented in the open state (low FRET, *E* = ~0.3) when they appeared on DNA. Fully complementarity between sgRNA and DNA drove Cas9_HNH1_ to quickly transit from the open state (lifetime ~ 100 ms) to and mainly stay at the intermediate state (medium FRET, *E* = ~0.6), whereas mismatches failed to cause such transition (Fig. 3a, S6b and c). Similar results were captured using Cas9HNH2 (Fig. S7). More strikingly, individual smFRET traces and transition density plots exhibited that conformation of Cas9_HNH1_ was highly plasticity and frequently transited to a high FRET state (*E* = ~0.9, Fig. 3b and S6a) only when mismatches between sgRNA and DNA at the PAM-distal end were 3 or less. 4 or more mismatches or nuclease dead Cas9 (dCas9) on perfectly matched DNA rarely sampled the high FRET state (Fig. 3 and S6d-e). Such strong correlation between sampling the high FRET state and Cas9 nuclease activity further supports that the high FRET state (*E* = ~0.9) captured in pre-steady state is the cleavage-competent state. Transient sampling the cleavage-competent state is sufficient for Cas9 to trigger the nuclease activity.

**Fig. 3.**
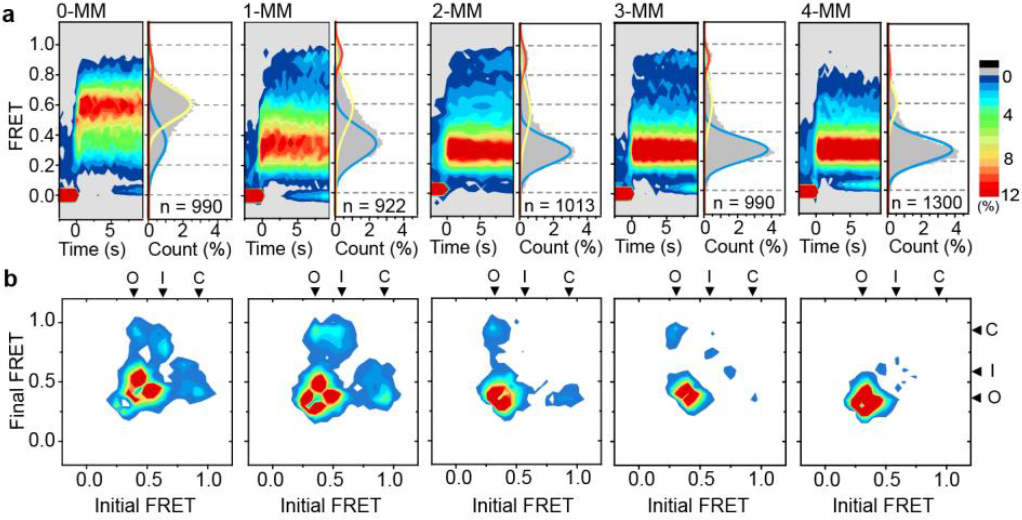
Dynamics of Cas9_HNH1_ captured in pre-steady state upon DNA binding. **a**, Time-dependent FRET contour plots (**a**, left of each sub-figure) reflecting evolution of FRET efficiency of Cas9_HNH1_ as the function of time upon binding to 0 – 4-MM DNA substrates. All movies were recorded at 500 ms per frame for 10 min and started 10 s prior to injecting labeled Cas9/sgRNA complexes. All smFRET trajectories were aligned to the first appearance of nonzero FRET states as *t* = 0 to construct time-dependent contour plots. Color codes are the same as Fig. 1c. Population histograms (**a**, right of each sub-figure) representing accumulative FRET efficiency distributions of the first 20 frames (red, yellow and blue lines represent closed, intermediate, and open states, respectively). **b**, Corresponding transition density plots showing the frequency of transitions between states. Each single molecule transition event of its initial and final FRET value was accumulated into 2D plot. Strikingly, Cas9HNH1 on 0 – 3-MM samples the closed state while Cas9_HNH1_ on 4 or more mismatches or dCas9 on full complementary DNA (Fig. S6d and e) rarely sample the closed state.

### Cas9 variants with enhanced or reduced specificity

Previous studies suspected that nuclease activity of Cas9 is energy-driven and ternary complex of dsDNA/sgRNA/Cas9 might possess more energy than it required for optimal specificity, which causes unwanted off-target cleavage (Fu et al., 2014). Based on such speculation, Cas9 variants, such as eSpCas9(1.1) (K848A/ K1003A/ R1060A) and Cas9-HF1(N497A /R661A /Q695A /Q926A), have been engineered to reduce interactions between Cas9 and DNA to achieve better specificities, whereas Cas9(L847R) containing a mutation to strengthen Cas9-DNA interactions displayed a reduced specificity (Kleinstiver et al., 2016; Slaymaker et al., 2016).

We introduced these mutants into Cas9_HNH1_ to examine how they affect cleavage activities (Fig. S8) and conformational dynamics of Cas9 (Fig.4 a-b and S9, Table S4). eSpCas9(1.1)_HNH1_ displayed slower cleavage rates and better specificity to DNA substrates than Cas9_HNH1_. Cleavage rate of Cas9(L847R)_HNH1_ towards 0-MM was slower than Cas9_HNH1_, however, it cleavage rate towards 4-MM was significantly faster than Cas9_HNH1_, which indicated that Cas9(L847R)_HNH1_ displayed a reduced specificity as discovered by previous study (Slaymaker et al., 2016). Thermodynamic and dynamic properties of Cas9-HF1_HNH1_ and Cas9(L847R)_HNH1_ both suggested that high fidelity is correlated with unstable closed conformational states (Fig. 4a) and slow transition rates towards the closed state (Fig. 4b), and *vice versa*. However, eSpCas9(1.1)_HNH1_ presented as an exception here, which has similar free energies as Cas9_HNH1_, whereas its effective transition rates to the closed state are 1.5-5 folds slower. Together, these results indicated that energy barriers of transit pathways towards the closed Cas9 state, which determine transition rates to the closed state, play more important roles than the thermodynamic stability (indicated by free energy) of the closed state. These energy barriers are modulated by interactions between Cas9 and dsDNA, including both target and non-target strands, as eSpCas9(1.1) and Cas9-HF1 were designed to weaken interactions of Cas9 with target strand and non-target strand, respectively (Kleinstiver et al., 2016; Slaymaker et al., 2016).

**Fig. 4.**
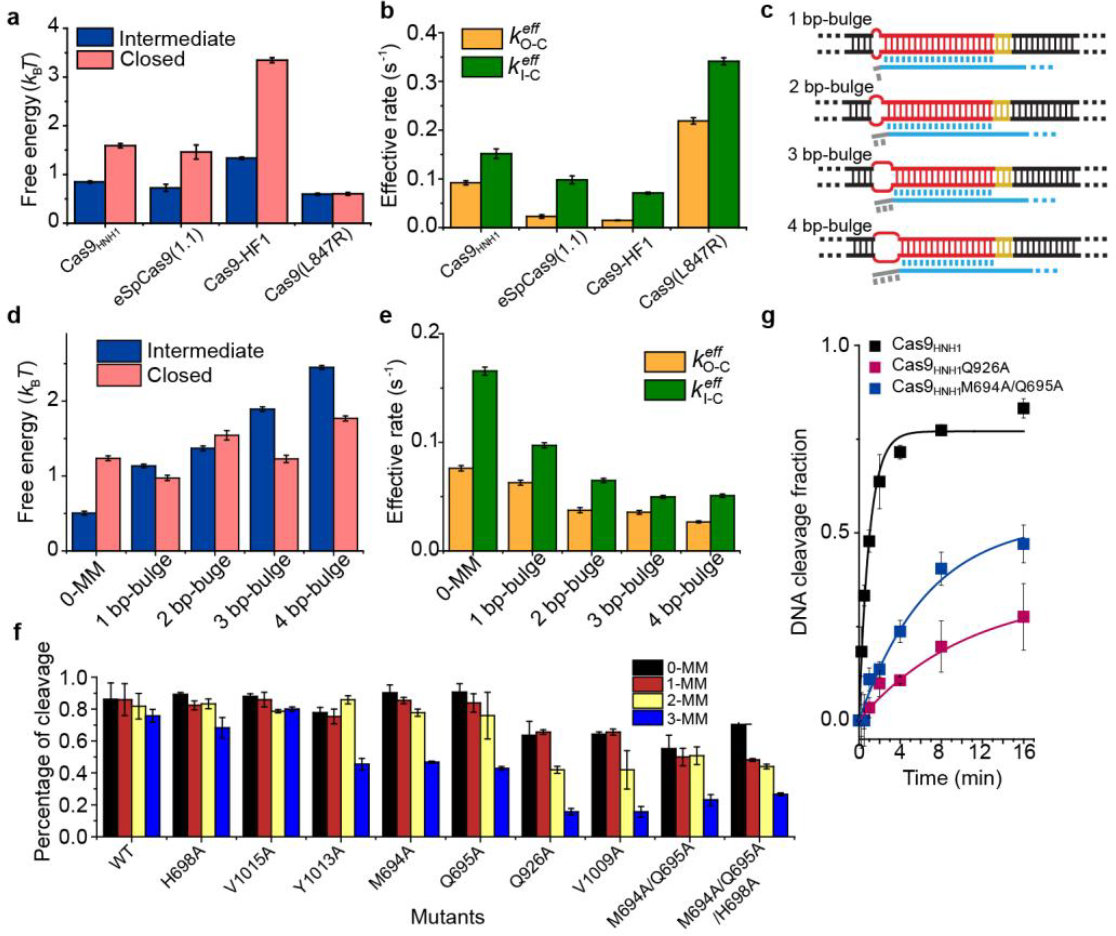
Conformational dynamics and cleavage activities of Cas9_HNH1_ modulated by its interactions with nucleic acids. **a-b**, Free energies of the intermediate and closed states (**a**) and effective transition rates from open and intermediate states to the closed states (**b**) of Cas9_HNH1_ and its variants. Specificity of Cas9 strongly correlates with the transition rates towards the closed state. **c**, Bulged DNA substrates design. Target DNA sequences are shown in red with the PAM in yellow. 1-4 internal mismatches on double-stranded DNA are shown as bubble. Matched and mismatched bases of sgRNA respond to target DNA are colored in blue and grey, respectively. **d-e**, Free energies of the intermediate and closed states (**d**) and effective transition rates (**e**) of Cas9_HNH1_ on 1-4bp bulged DNA substrates. Mismatches between DNA cannot fully compensate the penalties on stability and access rate of the closed state caused by mismatches between sgRNA and DNA. **f**, *In vitro* plasmid DNA cleavage assay of Cas9 variants for 30 minutes. **g**, Time-dependent *in vitro* plasmid DNA cleavage assay mediated by Cas9 variants. Percentage of cleaved DNA was quantified from agarose electrophoresis imaging and fitted by single exponential curves. Apparent cleavage rates of Cas9_HNH1_, Cas9_HNH1_Q926A, and Cas9_HNH1_M694A/Q695A towards fully-matched plasmids were 0.8±0.1, 0.10±0.04, and 0.15±0.04 min^-1^, respectively.

### Interactions between Cas9 and heteroduplex at the PAM-distal end

So far, we have clearly shown that introducing energy penalty during formation of dsDNA/sgRNA/Cas9 complex by creating mismatches between sgRNA and DNA or mutations on Cas9 can destabilize the closed Cas9 state and increase energy barriers towards the closed state to modulate its nuclease activity and specificity. We next introduced internal mismatches in dsDNA at the same positions that contained mismatches to sgRNA at the PAM-distal end (1~4-bp bulged DNA)(Fig. 4c and Table S1). Interestingly, in our smFRET assays, Cas9 on 1~4-bp bulged DNA showed different properties from that on perfect substrate (Fig. 4d-e and Table S2); similarly, the Doudna group have shown that *in vitro* cleavage rates of 4-bp bulged DNAs were 10-100 folds slower than that of a perfectly matched one (Sternberg et al., 2015). Together, these results indicated that energetic advantages gained from unwinding bulged-DNA duplex cannot fully compensate the penalties caused by mismatches between sgRNA and DNA. As a result, we hypothesized that Cas9 could proofread the formation of RNA/DNA heteroduplex at the PAM-distal end through interaction networks among them. Based on published structures (Jiang et al., 2016; Nishimasu et al., 2014), we introduced mutations on Cas9 residues which are likely to interact with heteroduplex at the PAM-distal end, and measured cleavage abilities of these Cas9 mutants (Fig. 4f-g). Our results showed that Cas9 containing even a single mutation at sites 694, 695, 926, 1009, or 1013 displayed attenuated cleavage activities, suggesting that these residues are likely involved in the interaction networks sensing PAM-distal end heteroduplex formation.

We introduced mutations of Q926A, V1009A, Y1013A, or M694A/Q695A into Cas9_HNH1_ to examine how they affect conformational dynamics of Cas9 (Fig. S10 and Table S4). We found that DNA binding was not affected by these mutations, because dissociation of all these Cas9_HNH1_ variants from dsDNA target was neglectable over tens of minutes. Comparing with Cas9_HNH1_, variants containing V1009A or M694A/Q695A displayed slower transition rates towards the closed state (Fig. S10b), and the close state of M694A/Q695A variant was less stable (Fig. S10a). Properties of other variants, including Q926A and V1009A, were similar as the ones of Cas9_HNH1_. Our discoveries suggested that mutation of a single residue is less likely to significantly alter conformational dynamics of Cas9 and to significantly improve its specificity. Therefore, Cas9 variants with enhanced fidelity usually contain several mutated residues (Kleinstiver et al., 2016; Slaymaker et al., 2016).

## Discussion

### Dynamic actions of Cas9

Cas9 mediated DNA recognition and cleavage is a highly dynamic multi-step process, including sgRNA binding, target searching, RNA strand invasion, R-loop expansion, and DNA cleavage (Gorski et al., 2017; Jiang and Doudna, 2017). The main focus of this study is how conformational dynamics of Cas9 govern its DNA cleavage after Cas9 stably binds on DNA target.

Several different structural conformations of Cas9 have been visualized by structural studies (Anders et al., 2014; Jiang et al., 2016; Jiang et al., 2015; Jinek et al., 2014; Nishimasu et al., 2014). Here, we directly captured that Cas9 spontaneously transits among three major global conformational states, even when it binds on fully complementary target DNA, which reflects significant dynamic motions of the catalytic HNH domain. Local conformation of the HNH domain can be further adjusted by formation of sgRNA/DNA heteroduplex at the PAM-distal end through a long-range allosteric communication. The local conformational flexibility of HNH domain has also been suggested by molecular dynamic simulations (Palermo et al., 2016) In addition, previous smFRET studies have revealed that fully-matched sgRNA/DNA heteroduplex also spontaneously transits between a zipped and an unzipped conformations on its PAM-distal end (Lim et al., 2016). Together, these results unambiguously demonstrate the highly conformational dynamic nature of dsDNA/sgRNA/Cas9 complex in both global and local scale.

While we were submitting our manuscript, two similar studies were deposited onto bioRxiv (Dagdas et al., 2017; Osuka et al., 2017). Using smFRET approach to capture dynamic motions of the HNH domain, both studies revealed that Cas9 exhibits three major conformational states, which is consistent with the open, intermediate, and closed Cas9 states we discovered. Interesting and puzzling, Dagdas et al. showed significant changes in smFRET distributions of Cas9 upon binding to various DNA targets, which is in line with ensemble FRET measurements from Sternberg et al. (Sternberg et al., 2014). On the other hand, Osuka et al. and our studies made similar observation that Cas9 shows similar smFRET distributions in its apo, sgRNA-bound, and dsDNA/sgRNA-bound forms. Agreeing with our smFRET measurements, ensemble FRET of Cas9_HNH1_ displayed minor changes upon binding to sgRNA and various DNA targets under our experimental conditions (Fig. S11 a and c). To address the discrepancies among these four studies, we tested many experimental conditions, including different temperature, reaction time, and buffer condition. We noticed that, heparin, a highly negative charge polymer which is likely to strongly interact with positively charged region of Cas9, was included in buffers by Dagdas et al. and Sternberg et al., but was not used by Osuka et al. and us. Our ensemble FRET experiments showed that, in the presence of heparin, Cas9_HNH1_ bound on targets containing 0-4 mismatches exhibited slightly larger FRET changes (Fig. S11 b and c). However, the FRET changes we captured were still smaller than what Sternberg et al. showed. To fully resolve the discrepancies, further studies are needed to understand how degree of polymerization and polymer size distribution of heparin affect its interactions with Cas9.

### A mechanism of high fidelity

The closed conformational state of Cas9 is least populated under all circumstances. Transient stay in the cleavage-competent closed state, in which the catalytic sites are in the correct positions towards their target sites, is sufficient for Cas9 to cleave its target DNA stands. Similar molecular behaviors have been discovered and proposed to be the molecular mechanisms to achieve high fidelity during aminoacyl-tRNA selection in elongation and DNA replication, during which ribosome/aminoacyl-tRNA/elongation factor Tu complex and DNA polymerase have been shown to fluctuate between non-active and active states, respectively, even in the presence of correct substrates (Geggier et al., 2010; Hohlbein et al., 2013; Lee et al., 2007). We suggested that Cas9 adopts a similar molecular mechanism to achieve its high specificity. The stability of the cleavage-competent closed state is evolved to be less stable than other non-active states. Therefore, only correct DNA target can trigger transient sampling towards the cleavage-competent state to complete cleavage, whereas DNA target containing mismatches is less likely or completely fails to cause such transient sampling. The fact that weakening or strengthening interactions between Cas9 and DNA leads to enhanced or reduced Cas9 specificity (Kleinstiver et al., 2016; Slaymaker et al., 2016), accompanied by a decreased or increased population of the cleavage-competent state (Fig. 4a and Table S4), respectively, further supports our proposed mechanism. In summary, well-tuned conformational dynamics between non-active and active states is the underlying molecular mechanism of Cas9 to achieve its high specificity.

### A proofreading mechanism

We proposed a kinetic scheme to conclude our findings (Fig. 5). Global structural conformation of Cas9 is highly dynamic and it spontaneously fluctuates among three major conformational states, mainly reflecting the conformational dynamics of HNH domain among the open, intermediate, and closed states. Due to a long-range allosteric communication between the HNH domain and the PAM-distal end region, RNA/DNA heteroduplex containing 3 or less mismatches triggers local motions of the HNH domain to correctly position its catalytic site to form a “cleavage-competent” closed Cas9 state; whereas 4 or more mismatches cause a similar but incorrect local position of the HNH domain that leads to a “cleavage-impaired” closed Cas9 state. We hypothesized that Cas9 presents a unique proofreading mechanism to ensure complementarity between sgRNA and DNA by sensing zipped heteroduplex (<= 3 mismatches) at the PAM-distal end, after binding of Cas9 on DNA is stabilized by PAM recognition and heteroduplex formation adjacent to PAM. Interactions between Cas9 and zipped heteroduplex at the PAM-distal end allow Cas9 to pass through the proofreading step, the last checkpoint, trigger transient formation of the “cleavage-competent” closed state, and permit DNA cleavage. It is likely to design new Cas9 variants with enhanced specificity, especially towards the PAM-distal end, by modifying residues involved in the proofreading step. In summary, our findings demonstrated that multi-level of global and local conformational dynamics of Cas9 interplay with each other to govern its nuclease activity and specificity.

**Fig. 5.**
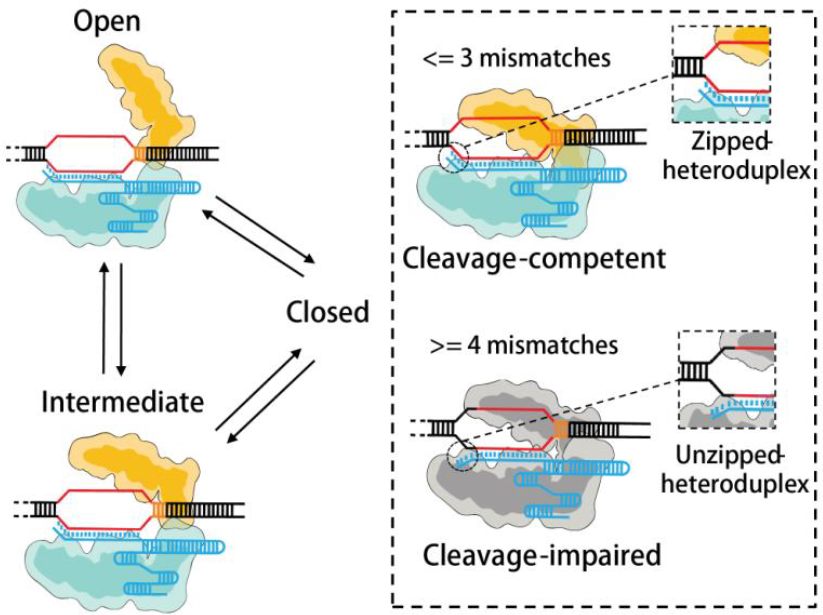
A proposed reaction scheme demonstrating how conformational dynamics of Cas9 governs DNA cleavage. HNH domain and α-helical lobe of Cas9 are shown in wheat and cyan, respectively. Target DNA sequences are shown in red with the PAM in yellow, and sgRNA is colored in blue. Cas9 spontaneously fluctuates among three major conformational states, mainly reflecting the significant conformational dynamics of the catalytic HNH domain. 3 or less mismatches between RNA/DNA heteroduplex trigger the HNH domain to correctly position its catalytic site displaying as a “cleavage-competent” closed Cas9 state. However, 4 or more mismatches lead to a “cleavage-impaired” closed Cas9 state. Taken together, we proposed that Cas9 presents a unique proofreading mechanism, to ensure complementarity between sgRNA and DNA by sensing formation of base pairs at PAM-distal end. Interactions between Cas9 and a zipped heteroduplex (<=3 mismatches) allow Cas9 to pass through the proofreading step, trigger transient formation of the “cleavage-competent” closed state, and permit DNA cleavage.

## AUTHOR CONTRIBUTIONS

MY and CC designed the experiments; MY, SP, RS, JL, and NW prepared materials and reagents; MY, SP and CC performed experiments; MY and CC analyzed the data and wrote the paper.

## Acknowledgments

This project was supported by funds from the National Natural Science Foundation of China (31570754), Tsinghua-Peking Joint Center for Life Sciences and Beijing Advanced Innovation Center for Structural Biology to CC.

